# Structure of the conjugation surface exclusion protein TraT

**DOI:** 10.1101/2025.05.27.656304

**Authors:** Nicolas Chen, Alfredas Bukys, Camilla A.K. Lundgren, Justin C. Deme, Hafez El Sayyed, Achillefs N. Kapanidis, Susan M. Lea, Ben C. Berks

**Affiliations:** Department of Biochemistry, University of Oxford, Oxford, OX1 3QU, United Kingdom; Center for Structural Biology, Center for Cancer Research, National Cancer Institute, Frederick, MD 21702, United States of America; Biological Physics Research Group, Department of Physics, University of Oxford, Oxford OX1 3PU, United Kingdom; Kavli Institute for Nanoscience Discovery, University of Oxford, Sherrington Road, Oxford OX1 3QU, United Kingdom

## Abstract

Conjugal transfer of plasmids between bacteria is a major route for the spread of antimicrobial resistance. Many conjugative plasmids encode exclusion systems that inhibit redundant conjugation. In incompatibility group F (IncF) plasmids surface exclusion is mediated by the outer membrane protein TraT. Here we report the cryoEM structure of the TraT exclusion protein complex from the canonical F plasmid of *Escherichia coli*. TraT is a hollow homodecamer shaped like a chef’s hat. In contrast to most outer membrane proteins, TraT spans the outer membrane using transmembrane helices. We use a novel microscopy-based conjugation assay to probe the effects of directed mutagenesis on TraT. Our analysis provides no support for the idea that TraT has specific interactions with partner proteins. Instead, we infer that TraT is most likely to function by physical interference with conjugation.

## Introduction

Bacterial conjugation is a form of horizontal gene transfer in which genetic material, most commonly plasmid DNA, is transferred from a donor bacterium to a recipient bacterium through direct cell-to-cell contact^1,2^. Conjugation is a major route for the spread of antimicrobial resistance genes between bacterial pathogens^3^ and thus of major biomedical importance. The *Escherichia coli* F plasmid was the first conjugative plasmid to be identified^4^ and continues to serve as a model system for dissecting the molecular mechanisms underlying bacterial conjugation^1,5-7^.

Conjugation systems encode a type IV secretion system (T4SS) to mediate DNA transfer between donor and recipient cells^8-10^. In the F plasmid system, the T4SS assembles a long extracellular pilus^11^. This pilus normally acts as a retractable grappling hook to bring the donor and recipient into close contact ^12^ in a step called mating pair formation. However, under some circumstances the pilus can also serve as the conduit for DNA transfer, enabling mating at a distance^13,14^. During conjugation, one strand of the plasmid DNA is transferred through the T4SS into the recipient cell. Upon entry to the recipient cell the complementary DNA strand is synthesised and the conjugal DNA becomes fully functional resulting in the recipient cell becoming a new donor cell (transconjugant).

Many conjugative plasmids encode systems that prevent redundant plasmid transfer into cells that already contain the same or a closely related plasmid^15,16^, a phenomenon known as plasmid exclusion^17^. In the *E. coli* F plasmid and in other conjugative plasmids of the IncF incompatibility group, exclusion is mediated by TraS and TraT proteins^18,19^. TraS blocks DNA transfer following mating pair formation (‘entry exclusion’) whilst TraT acts at an earlier stage to inhibit mating pair formation (‘surface exclusion’).

TraT is an outer membrane (OM) lipoprotein^18,20-23^ that is exposed at the cell surface^24-26^. It is reported that TraT only excludes donors carrying the parental plasmid, but not closely related plasmids, and therefore shows plasmid specificity^27,28^. The mechanism by which TraT enables surface exclusion has not been established.

Intriguingly, TraT has been implicated in several functions beyond its role in conjugation. It has long been recognized as a factor conferring resistance to serum complement^25,29-33^, and has also been reported to protect against phagocytosis^34^ and some bacteriophages^35^. Even more remarkably, TraT homologues are coded on the chromosome of many human and animal pathogens, in many cases without any obvious association with conjugation^15,25^ (and see also a detailed recent phylogenetic analysis^36^). Taken together these observations raise the possibility that TraT may function through a mechanism that is independent of specific interactions with the conjugation machinery.

In this work, we provide further insight into the mechanism of conjugal surface exclusion through structural characterization of the F plasmid TraT protein complex.

## Results

### Structure of the F plasmid TraT complex

To gain insight into the mechanism of surface exclusion we determined the structure of TraT from the F plasmid. Overproduced, affinity-tagged TraT was solubilised using the detergent dodecylmaltoside (DDM) and purified (Extended Data Fig. 1a,b). Cryo-electron microscopy (cryo-EM) of the TraT preparation revealed homogeneous ∼100 Å wide particles (Extended Data Fig. 1c) from which we were able to solve a 2 Å resolution structure of the TraT complex (Fig. 1a, Extended Data Fig. 2, Table S1).

**Figure 1.**
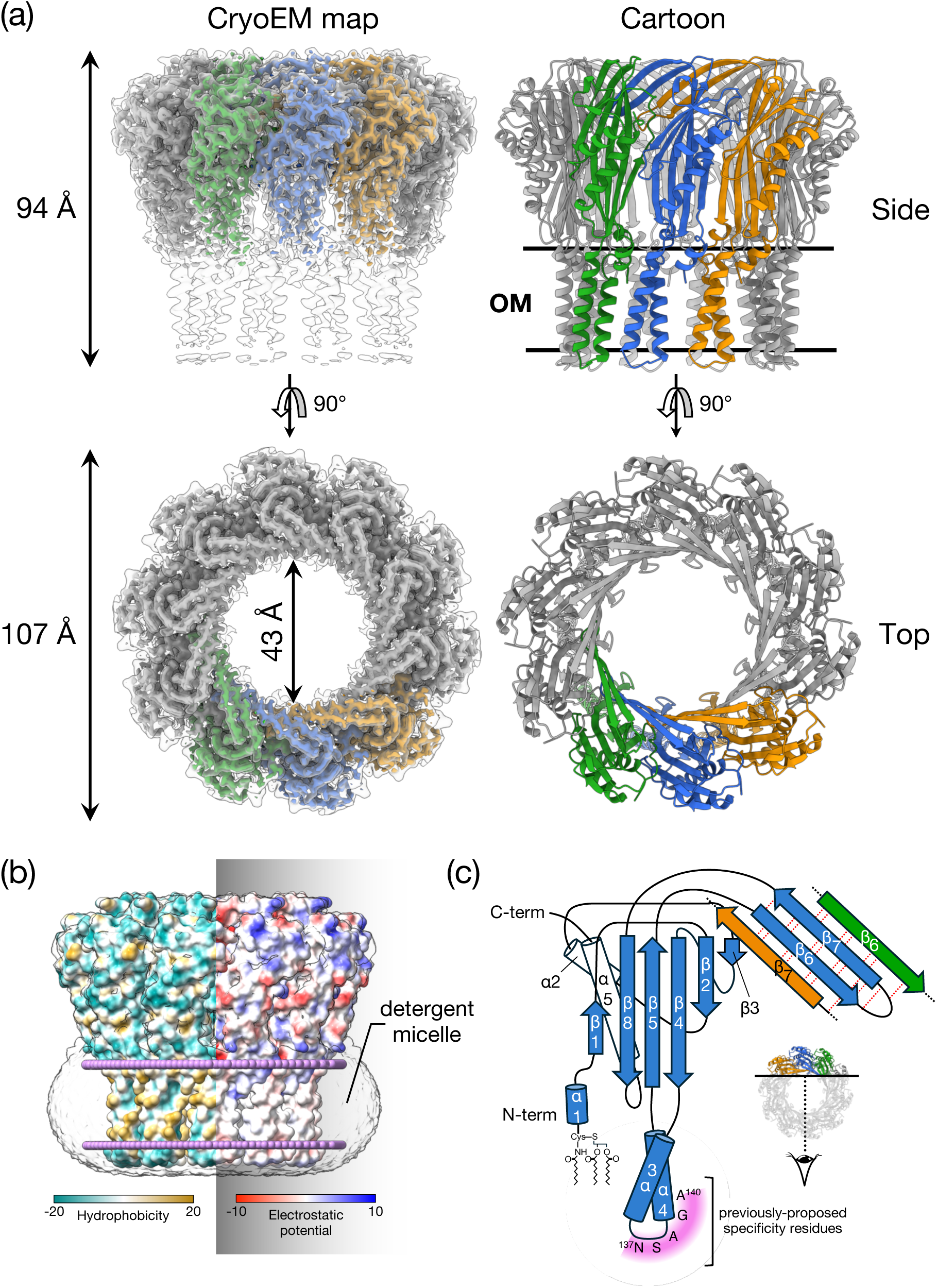
TraT forms a tubular homodecamer. **(a)** Cryo-EM structure of the TraT complex. The left hand panels overlay the unsharpened (transparent) and sharpened (solid colors and dark gray) cryoEM maps. The cartoon model of TraT is shown on the right. Three adjacent subunits within the TraT decamer are coloured. The position of the OM is shown based on the analysis in (b). **(b)** Surface hydrophobicity (left) and electrostatic potential (right) of the TraT model calculated using ChimeraX^60^. The position of TraT within the membrane (pink beaded lines) was predicted using the PPM server^66^ and by overlaying the cryoEM density for the DDM micelle. **(c)** Topology diagram of a TraT subunit viewed from inside the complex. The hairpin formed by strands β6 and β7 forms a continuous β sheet with the equivalent strands of the adjacent subunits (tan and green strands are from adjacent subunits as shown in the inset, hydrogen bonds in this sheet are represented by red dotted lines). Purple shading highlights previously proposed specificity-determining residues^27,28^.

TraT forms a homodecamer (Fig. 1a). The overall structure resembles a chef’s hat in shape but with an internal cavity extending along the entire long axis. The bottom of the ‘hat’ inserts into the OM bilayer as judged from the position of the detergent micelle and from the local surface hydrophobicity (Fig. 1b). The top of the ‘hat’ extends from the membrane, ending in a flattened top. Based on earlier biochemical experiments ^24,26^ the head is assigned to be on the extracellular face of the OM. The entire TraT polypeptide could be modelled but the density for the N-terminal lipid anchor is not resolved. Nevertheless, the N-terminal cysteine to which the lipid moiety is attached is appropriately positioned to allow the lipid group to be inserted into the membrane surrounding the complex.

The TraT monomer comprises three distinct elements: a globular core domain, two transmembrane helices (TMHs), and a long β-hairpin. The globular core domain is composed of two helices packed against a predominantly antiparallel five stranded β-sheet from which the TMHs and the β-hairpin extend (Fig. 1c). This core domain fold can also be found in a number of proteins that bind anionic oligosaccharides, with the closest structural homologue being LptE, a protein involved in lipopolysaccharide (LPS) transport^37^ (RMSD = 2.5Å). However, a functional role for TraT in LPS binding is unlikely because the residues essential for LPS binding in LptE^37^ are not conserved in TraT and there is no density in our structure that could correspond to bound LPS.

The membrane spanning portion of each TraT subunit extends from the core domain and consists of two TMHs connected by a short, tight loop of four amino acids (S136-A139). This loop does not extend significantly beyond the surrounding detergent micelle. Notably the TMHs from neighbouring TraT subunits do not pack tightly against each other, rendering them relatively mobile and poorly resolved.

The long β-hairpin (residues 180-205) projects from the membrane-distal end of the core domain. The hairpin structures of adjacent TraT protomers associate into a continuous β-sheet that forms a ring around the interior face of the open end of the TraT cavity (Figs. 1a,b).

In the native cellular environment the transmembrane end of the TraT cavity is almost certainly sealed by a lipid plug and so does not form an open channel across the outer membrane. We reach this conclusion because the transmembrane end of TraT has a hydrophobic interior, is filled with detergent density in the experimental structure, and is connected to the surrounding OM by gaps between the TMHs of adjacent subunits (Fig. 2). The most constricted regions of the TraT cavity are located at the β-sheet ring (43 Å diameter) and at the ‘top’ of the lipid plug (33 Å diameter)(Figs. 1a, 2).

**Figure 2.**
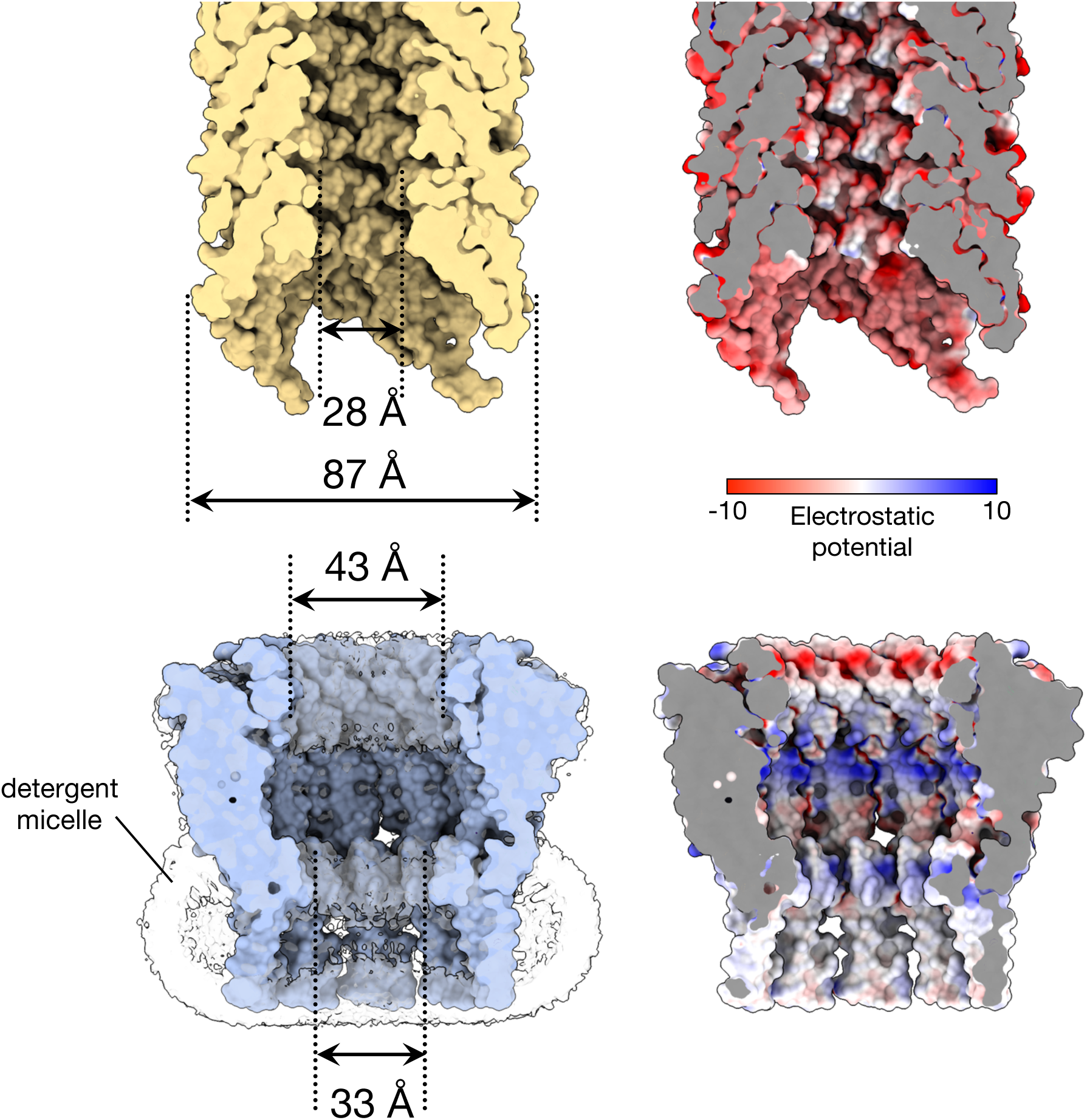
Comparison of the size and internal surface charge of TraT and the conjugation pilus. Shown are cut-throughs of the space filled structures of the pilus (top) and TraT complex (bottom, overlaid on the left with the cryoEM density for the detergent micelle). Surface electrostatic potential (right) was calculated using ChimeraX^60^. The pilus structure shown is that of the conjugation apparatus from the F-like plasmid pED208 and includes the phospholipid molecules that coat the internal face of the pilus tube.

It has previously been suggested that TraT might interact with the end of the conjugative pilus to prevent it from penetrating the OM^27^. To explore this possibility, we compared the TraT structure with that of the pilus shaft from the F-like plasmid pED208, a structure that we resolved at high resolution in this study (Fig. 2, Extended Data Fig. 3, Table S1). The diameter of the pilus shaft (87 Å) is similar to the diameter of TraT, suggesting possible size matching between the two structures. However, the entrance to the TraT cavity is only 43 Å in diameter and so too narrow for the pilus shaft to enter. A limitation of this comparison is that the pilus tip is likely made up of specialized proteins^27,38^ that are not present in our pilus shaft structure and which are likely to mediate any pilus interactions with TraT.

The cavity of TraT is markedly different from the lumen of the conjugative pilus through which DNA is transferred during conjugation (Fig. 2). As previously detailed^11^, the pilus lumen is narrow (28 Å diameter), lined with phospholipids, and negatively charged to reduce interactions with the moving DNA molecule. By contrast, the TraT cavity is much wider and has a non-uniform charge distribution, including both highly acidic and highly basic regions. These differences suggest that TraT is not structurally or electrostatically adapted for direct interaction with the transferring DNA.

### The previously proposed specificity region of TraT does not influence the exclusion phenotype

Plasmid exclusion systems are reported to only block conjugation from donors carrying the same plasmid^17^. In other words, exclusion systems exhibit specificity in their interactions with the conjugation machinery of the donor plasmid. In an earlier study of F-like plasmids it was reported that exclusion specificity is mediated in part by TraT^28^. In that study the authors identified the five amino acid sequence ^137^NSAGA^141^ in the F plasmid TraT complex (TraT_F_) as being responsible for this phenomenon. With the TraT structure now available this ‘exclusion specificity sequence’ can be seen to correspond to residues within the tight surface loop between the two TMHs of TraT_F_ together with the first two residues of the second TMH (Fig. 1c). In a later study, it was reported that TraT_F_ fails to exclude plasmid R100 even though it only differs from the sequence of TraT_R100_ at position 141^27^. Since residue 141 lies within the earlier-identified ‘exclusion specificity sequence’ this would appear to support the assigned role of this sequence in entry exclusion. However, the TraT_F_ sequence shown in ^27^ is incorrect and the TraT proteins from F and R100 are actually identical in sequence, leaving the difference in their exclusion effects unexplained. Given these uncertainties in the literature, we reinvestigated the effects of sequence changes in the proposed ‘exclusion specificity sequence’ of TraT. We compared three variations of the TraT specificity sequence: NSAGA, which is found in TraT from the plasmids F, R100, R6-5, and R1; NSAGG which is the incorrect F TraT sequence in ^27^; and SSAGA which is found in TraT from the plasmids pED208 and ColB K98.

To facilitate these experiments we developed a novel fluorescence microscopy-based conjugation assay which allowed more rapid data collection than traditional plating assays (Fig. 3). The method makes use of a fluorescence repressor operator system (FROS) ^39,40^ to identify conjugal plasmid transfer. In our assay the donor strain contains a F plasmid marked with a tandem array of *tetO* sites (F*_tetO_*) and the recipient strain expresses TetR-mNeonGreen (TetR-mNG). Conjugal transfer of the F*_tetO_* plasmid into the recipient strain results in TetR-mNG molecules binding to the *tetO* sites on the plasmid. This produces bright fluorescent foci which are automatically detected through subsequent image processing. To allow unambiguous identification of the donor cells in these experiments, donor cells were also engineered to produce chromosomally encoded mScarlet-I.

**Figure 3.**
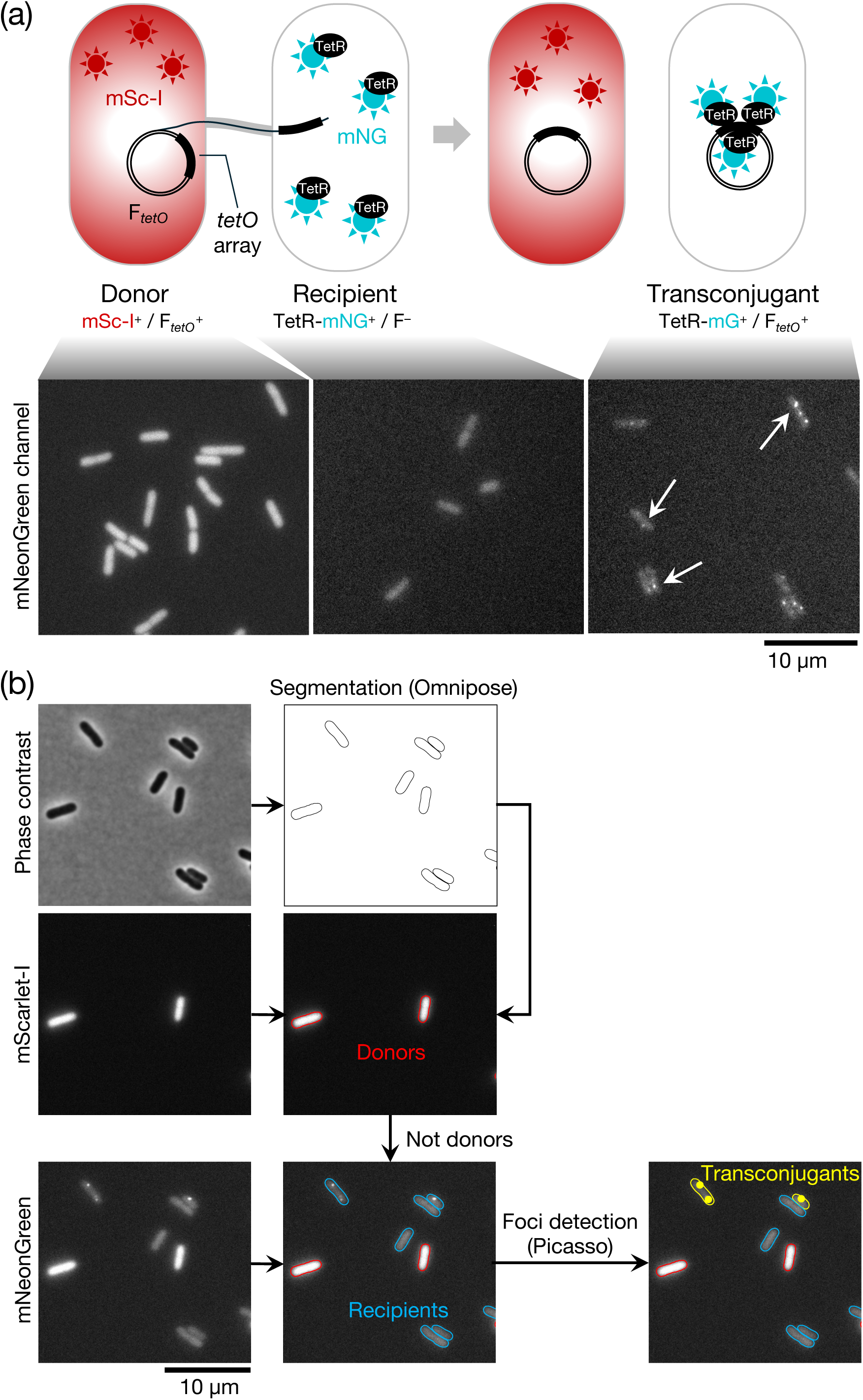
Microscopy-based conjugation assay. **(a)** (Top) Schematic overview of the conjugation assay (top). The donor carries a F plasmid bearing a *tetO* tandem array (F*_tetO_*^+^). The recipient produces a TetR-mNeonGreen fusion (TetR-mNG) that binds to the *tetO* array on the incoming F plasmid to form a fluorescent focus. The donor strain expresses cytoplasmic mScarlet-I (mSc-I) to allow the cells to be identified during fluorescence imaging. (Bottom) Representative fluorescence images in the mNeonGreen channel of the strains shown schematically above. The fluorescence of the donor marker (mScarlet-I) bleeds into the mNeonGreen channel so donor and recipient cells have to be separated by using the fluorescence in the mScarlet-I channel. **(b)** Microscopy data processing flowchart. The phase contrast image is segmented using Omnipose^62^ to extract cell outlines. Cells that display fluorescence in the mScarlet channel are labeled as donors, other cells are recipients. Fluorescent foci present in the mNeonGreen channel in recipient cells are detected using Picasso^63^ and cells with such foci assigned as transconjugants.

We examined the exclusion levels of recipient cells expressing TraT_F_ proteins bearing the three variations of the proposed ‘exclusion specificity sequence’ (Fig. 4a). We used immunoblotting to confirm that all three variants were produced at the same level (Fig. 4b). This analysis also showed that the recombinant TraT proteins were produced at concentrations close to those produced by the native F plasmid (Fig. 4b) and, therefore, that TraT function was being assessed at physiologically reasonable expression levels. In each case the TraT_F_ variant reduced conjugation efficiency (as measured by the mating ‘exclusion index’ (EI)) by the same amount (Fig. 4a). Thus, our data do not support the hypothesis that the proposed ‘exclusion specificity sequence’ results in TraT exclusion specificity.

**Figure 4.**
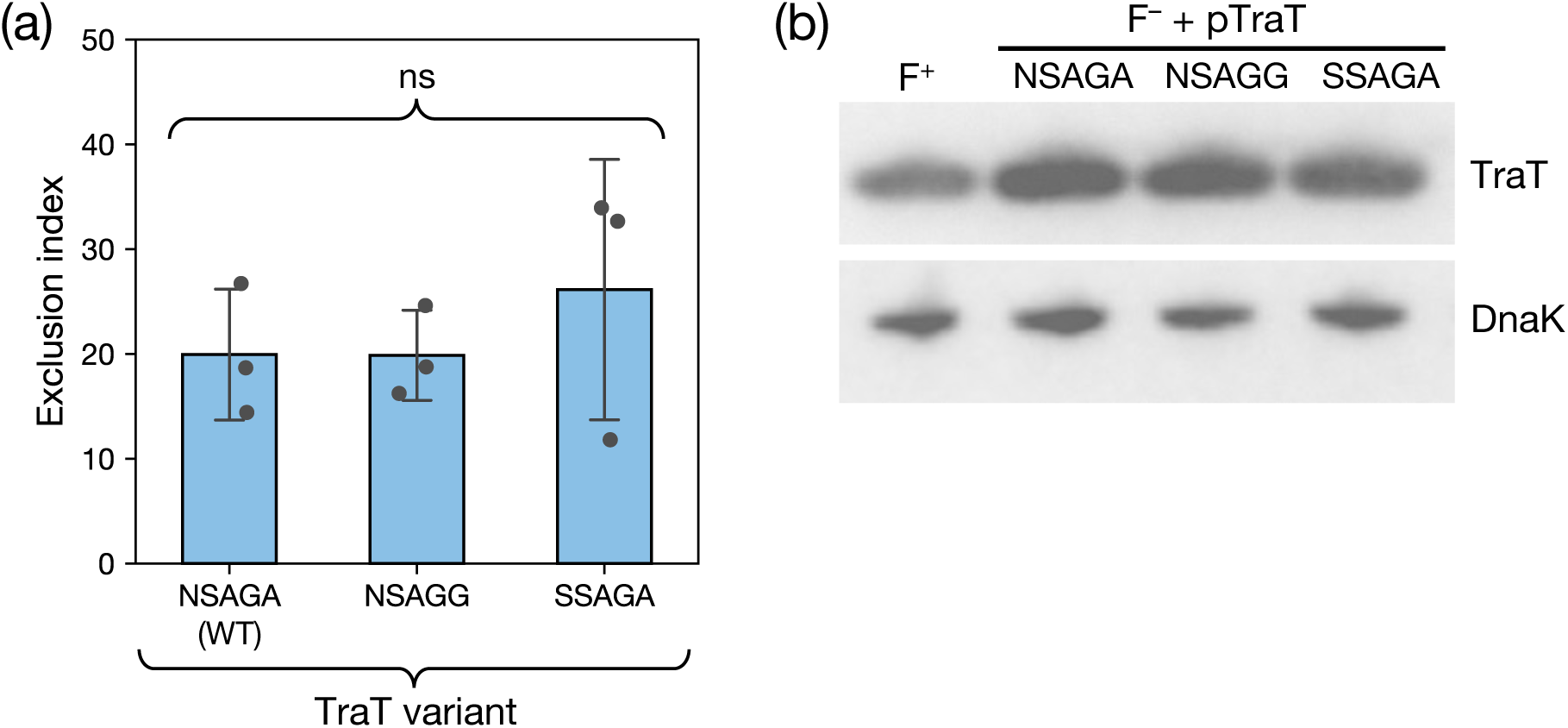
The previously proposed specificity residues of TraT do not affect surface exclusion. **(a)** Recipient cells contained a plasmid expressing the required TraT variant under the control of an IPTG-inducible promoter. The variants differ by the indicated sequence differences in the proposed specificity region encompassing residues 137 and 141 (WT, wild type sequence). Cultures were induced for 30 mins with IPTG prior to mixing donor and recipient cells (a) or immunoblotting (b). **(a)** Exclusion indices for the TraT variants determined using the conjugation assay described in Figure 3. Data bars represent the mean ± standard deviation obtained from 3 biological repeats (n=3), with each repeat represented as a point on the graph. A one-way F ANOVA was used to test the difference between the means of the three variants (ns, not significant at α=0.05). **(b)** Immunoblot comparing whole cell TraT levels in recipient strains expressing the indicated TraT variants (F^-^ + TraT lanes) or containing the F plasmid (F^+^ lane). DnaK is used as a loading control.

### Disruption of conserved residues on the internal surface of TraTF does not abolish surface exclusion

We calculated surface amino acid conservation for TraT_F_ in an attempt to identify residues that might interact with components of the donor conjugation apparatus or serve other important functional roles (Fig. 5a). This analysis revealed an invariant arginine residue (R213) on the internal surface of TraT. This residue appears unlikely to be required for structural stability of the complex and so is a strong candidate to have a functional role in TraT activity. The side chain of R213 is hydrogen bonded to a conserved aspartic acid (D170) on an adjacent β strand (Fig. 5a).

**Figure 5.**
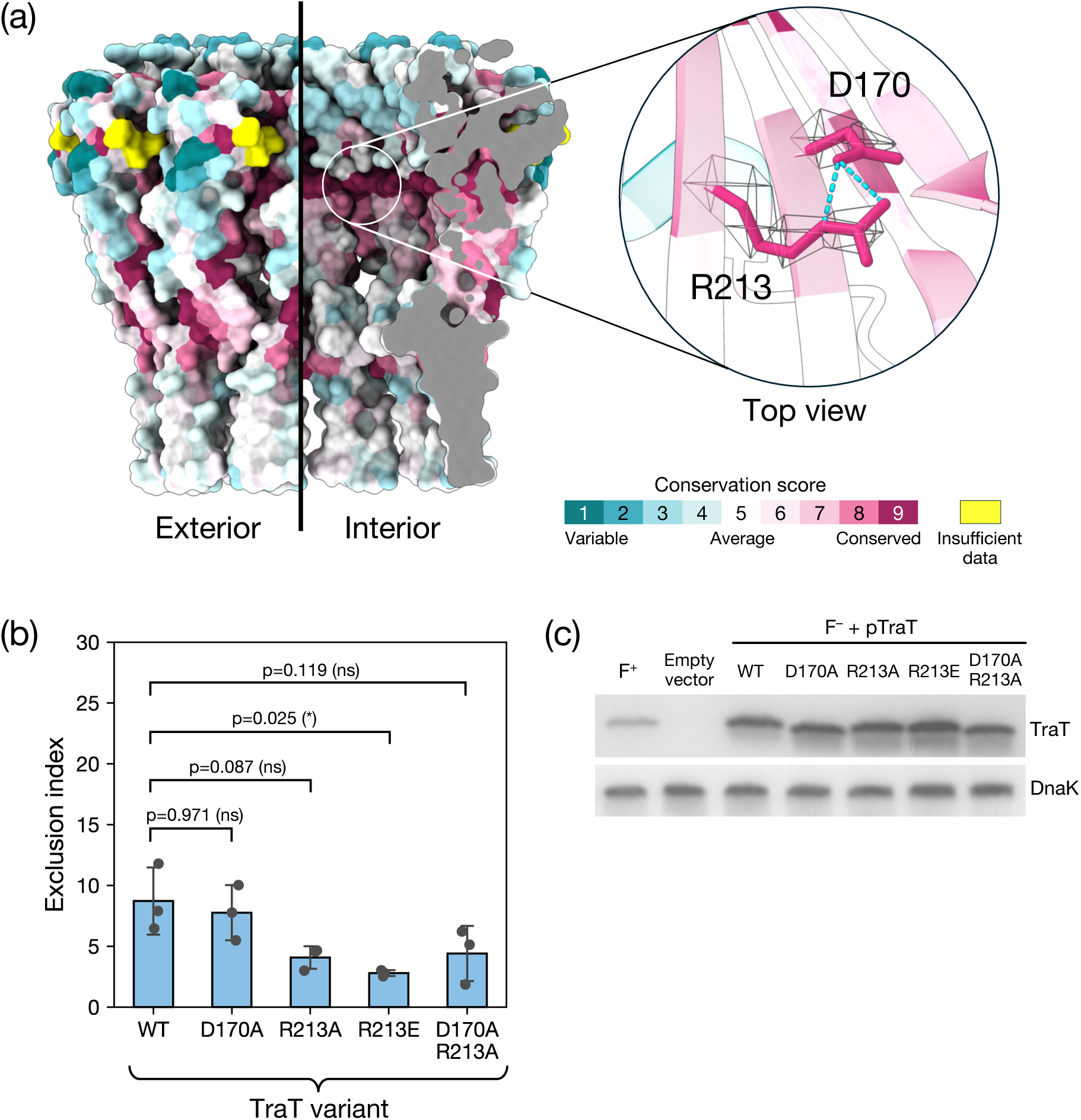
Disruption of two highly conserved residues in the TraT lumen does not abolish surface exclusion. **(a)** Surface conservation score calculated by Consurf^64^ mapped onto the structure of the TraT complex and viewed both from the exterior (left) and interior (right, cutaway view) of the complex. The inset shows a magnified view of a pair of highly conserved residues (R213 and D170) that interact with one another on the interior of the complex through side chain hydrogen bonds (cyan dashed lines). The EM density for the side chains of these residues is shown over the TraT model. **(b,c)** Analysis of TraT variants containing the indicated amino acid substitutions. Recipient cells contained a plasmid expressing the required TraT variant under the control of an IPTG-inducible promoter. Cultures were induced for 30 mins with IPTG prior to mixing donor and recipient cells (b) or immunoblotting (c). WT, wild type. **(b)** Exclusion indices determined using the conjugation assay described in Figure 3. Data bars represent the mean ± standard deviation obtained from 3 biological repeats (n=3), with each repeat represented as a point on the graph. A one-way F ANOVA was used to test the difference between the means of the three variants (significant at p_adj_=0.016). The variants were then analyzed by pairwise Tukey’s HSD tests relative to the WT. ns: non-significant (p_ajd_ ≥ 0.05), *: p_ajd_ < 0.05. **(c)** Immunoblot comparing whole cell TraT levels in recipient strains either expressing the indicated TraT variants (F^-^ + TraT lanes) or containing the empty expression plasmid (Empty Vector), or bearing the F plasmid (F^+^ lane). DnaK is used as a loading control

To assess the importance of R213 and D170 to TraT_F_ function we altered their side chain properties by either neutralizing their charge (D170A, R213A variants, or the combination of the two substitutions) or reversing it (R213E variant). We then performed conjugation assays to assess the impact of these substitutions on surface exclusion (Fig. 5b). A control experiment confirmed that the variant and wild type TraT proteins were expressed at the same levels (Fig. 5c). All the variant TraT proteins retained the ability to mediate surface exclusion (Fig. 5b). For the variants with a substitution of R213, the EI was reduced relative to the wild type protein. However, this change was only marginally significant for the R213E substitution (Tukey’s HSD test, p_adj_ =0.025) and not statistically significant for the other variants. We, therefore, conclude that D170 and R213 are not essential for TraT function.

### TraTF fails to form a complex with Complement Factor H

TraT_F_ is a resistance factor against serum complement^25,29-33^ but the molecular basis of this activity has been unclear. Recently it has been reported that TraT from the fish pathogen *Edwardsiella tarda* binds complement factor H (CFH) ^25^. We investigated whether this interaction with CFH could also be demonstrated for TraT_F_ as a route to structurally characterise the basis of complement resistance by TraT. However, we were unable to detect complex formation between mouse CFH and our purified TraT_F_ protein (Extended Data Fig. 4) and so could not confirm the reported interaction between TraT and CFH.

## Discussion

The structure of the TraT conjugation surface exclusion complex reveals a homodecameric assembly with no obvious structural counterparts among other characterized protein complexes. Each TraT subunit has a lipid anchor, a feature that is typically found on peripherally anchored OM proteins. However, the structural data show that TraT is actually an integral OM protein in which a large globular domain associates with the membrane bilayer through a pair of TMHs in each subunit (Fig. 1). TMHs are exceedingly rare in OM proteins due to the difficulty in preventing their insertion into the IM as they are trafficked to the OM. In this respect TraT is a highly unusual OM protein and a notable addition to the very small group of OM proteins that are known to contain transmembrane helical segments^41,42^.

The structure of the TraT complex does not immediately suggest the mechanism of surface exclusion. Nevertheless, the structural data provide new insights into the plausibility of different potential mechanisms^43^.

One possible mechanism for surface exclusion is that TraT intercepts the tip of the conjugative pilus, preventing it from penetrating the OM of the recipient cell. This mechanism would be consistent with the observation that TraT blocks induction of the envelope stress responses that take place during conjugation^16^. A pilus-interception mechanism implies specificity in the interaction between the pilus and TraT. Such an interaction would be expected to require a conserved interaction interface on TraT. However, our structural work reveals no substantive region of high sequence conservation on the surface of TraT (Fig. 5a) and our mutagenic analysis shows that the most conserved surface residues are not required for TraT function (Fig. 5b,c).

Alternatively, if TraT is responsible for the plasmid specificity of conjugation exclusion, as suggested by some^27,28^, but not all^18^, studies then this should be mediated through the proposed pilus-TraT contact. However, we were not able to substantiate the presence of the previously proposed ‘exclusion specificity sequence’ in TraT^28^. We speculate that the earlier studies may have measured differences between TraT variants because they did not control for TraT expression levels, which are known to influence TraT activity^18^.

Another potential mechanism of surface exclusion^18,44^ is that TraT shields receptors on the recipient cell surface from interacting with either the pilus or the donor surface protein TraN^45,46^ which stabilizes mating pair formation. In the specific case of the F plasmid, the receptor for TraN is the abundant eight-stranded OM barrel protein OmpA^45,47^. The transmembrane cavity of the TraT complex would be large enough to sequester a single copy of OmpA away from incoming TraN molecules. However, we did not find OmpA co-purifying with TraT. In addition, TraN proteins from other IncF plasmids interact with receptor proteins that are much larger than OmpA^45^ and these proteins could not be housed within the TraT cavity. Thus, the structure of TraT does not support a receptor-shielding model for surface exclusion. A receptor shielding mechanism is also disfavoured by the relative stoichiometries of TraT and OmpA, since OmpA monomers are in a ∼30-50 times molar excess in the OM over the decameric TraT^18^.

Conjugal TraT proteins are known to block surface processes other than conjugation, including attack by serum complement. Moreover, TraT family proteins are often found in genetic contexts unrelated to conjugation^43^. These observations suggests that TraT might not operate through a mechanism that specifically targets the pilus tip or other components involved in mating pair formation but may, instead, act through a more generic interference with cell surface processes. Based on this idea, we now suggest that TraT mediates surface exclusion by altering the physical properties of the OM in a way that reduces the efficiency of pilus tip insertion. This model would be consistent with the inhibition of complement killing by TraT because this process requires the insertion of pore-forming proteins into the OM^48^. It may also explain why very high levels of TraT are needed to mediate surface exclusion^18,24,49,50^. It is not obvious how the structure of TraT could achieve this proposed effect, although we note that the TraT complex is a highly rigid structure that is likely to strongly withstand external mechanical forces.

In summary, TraT forms a highly unusual and abundant OM complex that protrudes from the OM surface into the extracellular space. The structure of TraT provides a molecular framework with which to unravel the basis of surface exclusion and complement resistance.

While this manuscript was in preparation the structure of TraT was also reported by other authors^36^.

## Data availability

Cryo-EM density maps are available in the Electron Microscopy Data Bank (EMDB) with the following accession numbers: EMD-47469 and EMD-48768.

Atomic coordinates are available in the Protein Data Bank with the following accession number: PDB 9E2V and PDB 9MZT.

## Code availability

The source code for processing of microscopy data is available at https://github.com/alfbukys/conj_assay. The source code used to perform statistical tests is available at https://github.com/nypchen/TraT-microscopy.

## Acknowledgements

We thank Kevin Foster for access to the fluorescence microscope used in this study and Rachel Jones for statistics advice. We thank Isobel Bender for producing the TraT antiserum. This research was funded in part by Biotechnology and Biological Sciences Research Council grant BB/S007474/1, European Research Council Advanced Award 833713, and a Titular Clarendon Fund Scholarship in partnership with the Merton College Biochemistry Scholarship and Biochemistry Studentship to AB. This research was supported in part by the Intramural Research Program of the NIH.

## Author Contributions

NC carried out all experiments except the following. The fluorescence conjugation assay was developed by AB with assistance from HES and the conjugation experiments were then carried out and analyzed by AB and NC. CL, JCD, and SML collected electron microscopy data and determined all structures. The work was supervised by BCB, CL, SML, and AK. All authors interpreted data, produced figures, and revised the manuscript.

## Materials and methods

### Strain and plasmid generation

All strains and plasmids used in this study are listed in Supplementary Table S2. All non-conjugative plasmids in this study were generated using Gibson assembly using the primers in Supplementary Table S3. All plasmid constructs were verified by sequencing.

Plasmid pNC60 for the isolation of TraT was constructed by introducing the F plasmid *traT* gene into a pET22b-derived vector that fuses a MDIGINGT linker and Twin-Strep tag to the C-terminus of the inserted protein.

Plasmids to test the function of TraT variants in conjugation assays were constructed by first cloning *traT* into pQE80L (Qiagen) to produce pNC56 and then introducing the required substitutions using a KLD mutagenesis (NEB).

For conjugation experiments the F plasmid (pOX38-Kan^R^) was marked with a 22-repeat *tetO* array inserted between the *ygfA and ygeB* genes using lambda-red recombineering as described^51^ with the linear insertion fragment amplified from pKD3_tetO using primers #72AB and #73AB. The chloramphenicol resistance cassette was flipped out and the resulting plasmid F*_tetO_* plasmid was conjugated into MG1655 *mScarlet-I*::Tn*7* ^52^, generating the donor strain used in the conjugation assays. The recipient strain AB_2 contains a *tetR*-*mNeonGreen* fusion under the control of the native *rpoN* promoter and the *rrnB* T1 and T2 terminators. The reporter was inserted into the *E. coli* chromosome via scarless recombineering ^53^ using pAB5, disrupting the open reading frame of *lacZ* and fully deleting *lacI*.

### Purification of the TraT complex

An overnight culture of *E. coli* C41 ^54^ containing pNC60 was diluted 1:50 in LB medium ^55^ containing 100 µg/ml of ampicillin. The diluted culture was incubated at 37 °C until OD_600_ = 0.7, then expression of *traT* was induced with 1 mM IPTG and the cells cultured for a further 3.5 hrs. Cells were harvested by centrifugation at 7,500*g* for 20 min. The cell pellets were resuspended in lysis buffer (phosphate buffer saline tablets [1 tablet/200 ml] (Sigma-Aldrich), 400 μg/ml lysozyme (Fluka), 30 μg /ml DNAse I (Sigma-Aldrich), UltraCruz EDTA-free protease inhibitor cocktail tablets [1 tablet/50 ml buffer] (Santa Cruz Biotechnology)) at a ratio of 5 ml buffer per g of cell pellet. Cells were lysed by three passages through a French press at 10,000 psi. The lysate was clarified by centrifugation twice at 30,000g for 30 min at 4°C. Membranes were recovered from the supernatant by centrifugation at 200,000*g* for 1.5 hrs at 4°C. Each gram of membrane pellet was resuspended in 8 ml of membrane resuspension buffer (100 mM Tris, 150 mM NaCl, UltraCruz EDTA-free protease inhibitors [1 tablet/50 ml buffer], pH 8 at 10 °C). One milliliter aliquots of the resuspended membranes were snap-frozen in liquid nitrogen and kept at −80°C for further use. After thawing, a 10% (w/v) stock solution of n-dodecyl-β-D-maltoside (DDM) (Anatrace) was then added to a final concentration of 1% (w/v), and the mixture was incubated at 10°C for 2 hrs with gentle agitation to solubilize the membranes. The mixture was centrifuged at 200,000*g* for 30 min at 4 °C to remove insoluble material. The supernatant was amended with BioLock (IBA Lifesciences) at 0.5 ml per 7 ml of solubilized membranes and applied to a Streptactin XT 4Flow high capacity affinity purification column (IBA Lifesciences). The column was washed with 30 ml of wash buffer (100 mM Tris, 150 mM NaCl, 0.2% DDM, 1 mM EDTA, pH 8 at 10°C) and eluted with 6-12 ml of wash buffer containing 50 mM biotin. The eluate was concentrated 20-fold using a 10kDa molecular weight cut-off centrifugal concentrator (Merck Millipore), filtered using a 0.22 µm spin filter (Merck Millipore), and run through a Superose 6 increase 10/300GL (Cytiva) size exclusion chromatography column pre-equilibrated with 100 mM Tris, 150 mM NaCl, 0.02% DDM, pH 8 at room temperature. Fractions containing purified TraT were identified by SDS-PAGE followed by Coomassie staining.

### Pilus purification

The method was based on that outlined in^11^. One hundred microliters of an overnight culture of cells free of non-conjugative pili and flagella (MC4100-AN) carrying pED208 were plated on 120×120 mm LB agar plates, and incubated overnight at 37°C. The next day, cells were scraped off using an L-shaped spreader and ice-cold buffer (25 mM HEPES pH7.15, 150 mM NaCl) (5 ml buffer/plate). An extra 10 ml of buffer was used to rinse all plates to increase the yield. Pili were shaved off using by homogenizing ≈30 times using a Dounce homogenizer. Shaved cells were centrifuged at 10,000g for 20min at 4°C. The supernatant was centrifuged again to remove residual cells. The supernatant was concentrated ≈20 times using a centrifugal concentrator (100k MWCO) (Merck Millipore).

### TraT-CFH pulldown

100 μl of either purified TraT-TS (1.13 mg/ml) in binding buffer (100mM Tris, 150mM NaCl, 0.02% DDM, pH8) or binding buffer only were mixed with 1 µl of MagStrep beads (IBA Lifesciences, Cat No. 2-5090-002) that had been washed and equilibrated according to the manufacturer’s protocol. After 30 min of incubation with gentle agitation the beads were recovered and washed 2 times with 200 µl of binding buffer. 5 μl of mouse CFH (Complement Technology, Cat. No. A137) (1.05 mg/ml) were added to the beads and the mixture was incubated for another 30 min. The beads were recovered and washed 3 times with 200 µl of binding buffer. The beads were then incubated with 25 µl of binding buffer containing 50 mM biotin for 10 min before separation of the beads and eluted fraction. Elution was repeated 5 times, before the beads were boiled at 100°C for 5 min in Laemmli buffer (62.5 mM Tris pH 6.8, 0.02% SDS, 0.1 M DTT, 10% glycerol). Samples of the different steps were then analyzed by SDS-PAGE, followed by Coomassie staining.

### Antisera used

Polyclonal antibodies against the purified TraT complex were raised in rabbits. The following commercial antisera were used: anti-DnaK (ADI-SPA-880-D Enzo Lifesciences), anti-mouse IgG peroxidase conjugate (A4416 Merck), and anti-rabbit IgG peroxidase conjugate (31462 Pierce).

### Cryo-EM sample preparation, data collection and processing

Purified TraT (3 μL, 12 mg/ml) or pili (3 μL) samples were applied onto glow-discharged (30 s, 15 mA) holey carbon-coated grids (Quantifoil Au, 300-mesh, R1.2/1.3). The sample was adsorbed for 10 s before blotting. Grids were blotted for 3-6 s, at 4 °C with 100% relative humidity using a Vitrobot Mark IV (Thermo Fisher Scientific). Grids were subsequently plunge-frozen in liquid ethane cooled by liquid nitrogen.

Cryo-EM data for both samples were collected in counting mode in Electron Event Representation (EER) format on a cold field emission gun (CFEG) equipped Titan Krios G4 transmission electron microscope (Thermo Fisher Scientific) operated at 300 kV. Selectris X imaging filter (Thermo Fisher Scientific) with a slit width of 10 eV was used in conjunction with a Falcon 4i direct electron detector (Thermo Fisher Scientific). Data were collected with a physical pixel size of 0.732 Å and a total dose of ∼54 e^-^/Å^2^.

For TraT the initial preprocessing steps including patched motion correction (20 × 20), contrast transfer function (CTF) parameter estimation, particle picking, extraction, and reference-free 2D classification were performed using SIMPLE 3.0^56^. All subsequent processing was carried out in cryoSPARC 3.3.1^57^ as described in the workflow (Extended Data Fig. 2). The final global resolution was estimated as 2.01 Å using a 0.143 criteria and gold standard FSC. Data processing for the pilus sample (Extended Data Fig. 3) was entirely within cryoSPARC 3.3.1 with the exception of particle polishing performed in RELION^58^. EMD4046 was used as a starting volume for initial alignment with starting helical parameters of 28.1° twist, 12.5 Å rise which refined to 28.203° twist and 12.086 Å rise. Global resolution was estimated as 2.11 Å using the 0.143 criterion and gold standard FSC.

### CryoEM model building and validation

An AlphaFold2^59^ model of a TraT decamer was rigid-body fitted into the Cryo-EM density map using ChimeraX^60^. Subsequent manual model building was iteratively carried out in coot^60^ using both sharpened and unsharpened maps for enhanced interpretability. Models were refined using real-space refinement and symmetry restraints in PHENIX^61^ to generate the model described in Table S1. Validation of the refined model was performed using MolProbity^58^ to assess structural integrity and stereochemical quality, including evaluation of bond lengths, angles, torsion outliers, and Ramachandran statistics. Map-to-model correlation coefficients were also used to verify the accuracy of model fitting. For the pilus, pdb 5lfb was roughly positioned in the high-resolution volume and manually adjusted in coot. Refinement in PHENIX yielded the model described in Table S1.

### Microscopy conjugation assays

Overnight cultures of donor (MG_mSC containing F*_tetO_*) and recipient strains (AB_2 with the appropriate pNC56-derived *traT* expression plasmid) were each diluted 1:50 in LB medium, supplemented in the case of the recipient strains with 100 µg/ml of carbenicillin. The cultures were then grown to OD_600_ = 0.4-0.7 at which point the recipient culture was supplemented with 1 mM IPTG to induce expression of *traT* and cultured for a further 30 min. Recipient cells were washed 2 times with fresh LB to remove carbenicillin and both cell suspensions were adjusted to OD_600_ = 1.5. 400 μl of recipient cells were mixed with 80 µl of donor cells (a 1:5 donor:recipient ratio), and the volume was adjusted to 800 µl with fresh LB. The mixtures were incubated with shaking at 37°C for 30 min to allow conjugation. The cell mixtures were then spotted onto agarose pads and imaged on a Zeiss AxioObserver widefield microscope equipped with a Colibri 7 LED light source and using a Plan-Apochromat 63x/1.40 oil immersion lens with phase contrast. Images were captured on a Sony ICX 694, EXview HAD CCD II sensor. Images were captured for 150 ms using 20% intensity 555 nm LED light (to visualize mScarlet-I), for 100 ms using 100% intensity 475 nm LED light (to visualize mNeonGreen), and for 200 ms using 15% transmitted light intensity for phase contrast.

### Microscopy data processing and statistics

Fluorescence microscopy image stacks were processed in the BacSEG Napari plugin (https://github.com/piedrro/napari-bacseg) which implements image segmentation via Omnipose^62^ and single molecule localisation via Picasso^63^. We used the Omnipose ‘bact_phase_omni’ model to segment the cells in the phase contrast channel. Processed data were manually curated to remove erroneous segmentation caused by agarose pad imperfections and image focusing errors. Cell segmentation coordinates, transconjugant F plasmid localisation coordinates, and the donor mScarlet-I fluorescence images were used to calculate the conjugation efficiencies for each mating experiment using a custom Python script.

Conjugation efficiencies were calculated using the following formula:

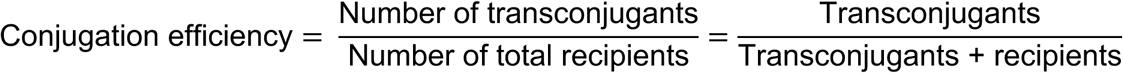

Exclusion indices for TraT-expressing recipients were calculated using the following formula:

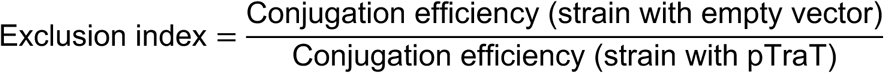

Data was collected from 3 distinct biological samples for each TraT variant (n=3). Statistical tests were performed using a one-way F ANOVA. Assumptions were verified using Shapiro’s (normality of residuals) and Bartlett’s (equality of variances) tests. When the ANOVA was significant (p<0.05), a Tukey’s Honestly Significant Difference (HSD) *post-hoc* test was performed for pairwise comparisons and to obtain adjusted p-values (p_adj_). The data from 45-50 fields of view per sample (10,000-100,000 cells) were combined for each biological repeat.

### Sequence conservation

The conservation score of TraT residues was calculated with the Consurf online server ^64^ using default parameters. Under these settings, the multiple sequence alignment is calculated from Uniref90 ^65^, a database in which sequences with >90% identity are clustered.

---

**Extended Data Figure 1.**
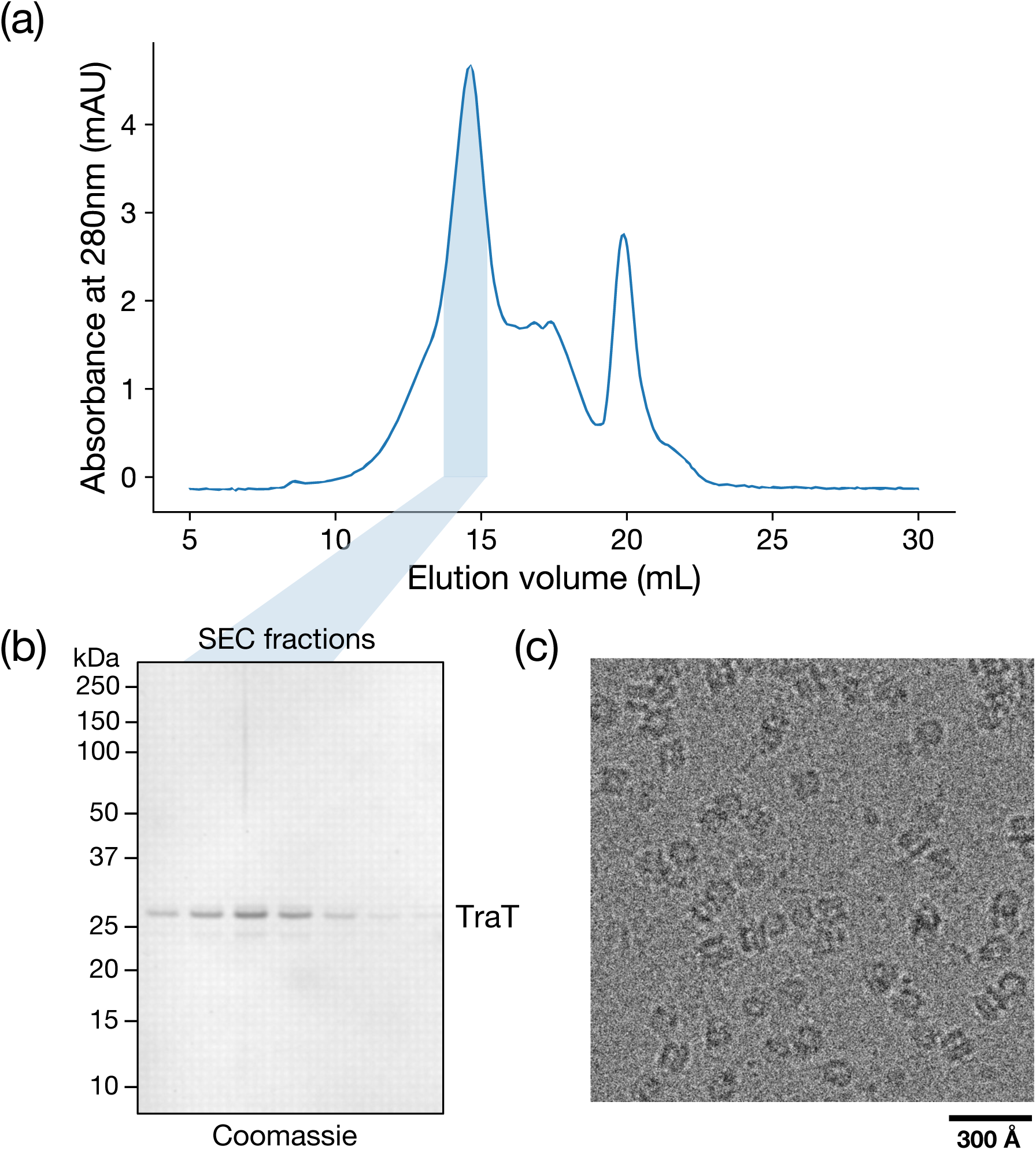
Purification of the F plasmid TraT complex. Twin-Strep tagged recombinant TraT was solubilized from membranes using DDM, and isolated by streptactin affinity chromatography followed by SEC. **a,b** SEC elution profile **(a)** and Coomassie stained SDS-PAGE gel **(b)** of the SEC peak fractions. **c,** Cryo-EM image of the pooled fractions shown shaded in blue in (a,b).

**Extended Data Figure 2.**
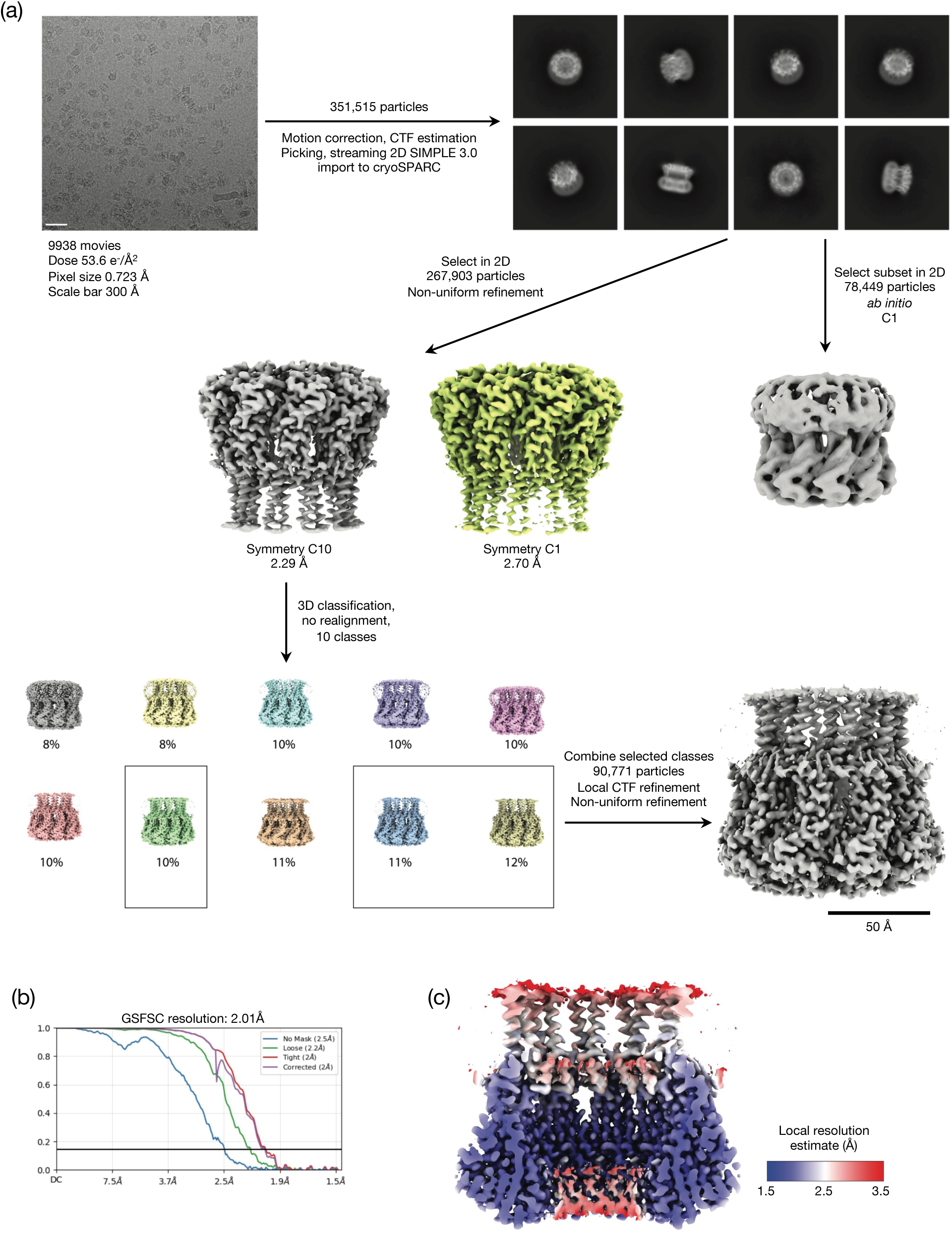
Cryo-EM workflow for the F plasmid TraT complex. **a**, Image processing workflow, Micrograph scale bar is 300 Å. **b**, Gold-standard Fourier Shell Correlation (FSC) curves used for global resolution estimation. **c**, Local resolution estimate of the volume.

**Extended Data Figure 3.**
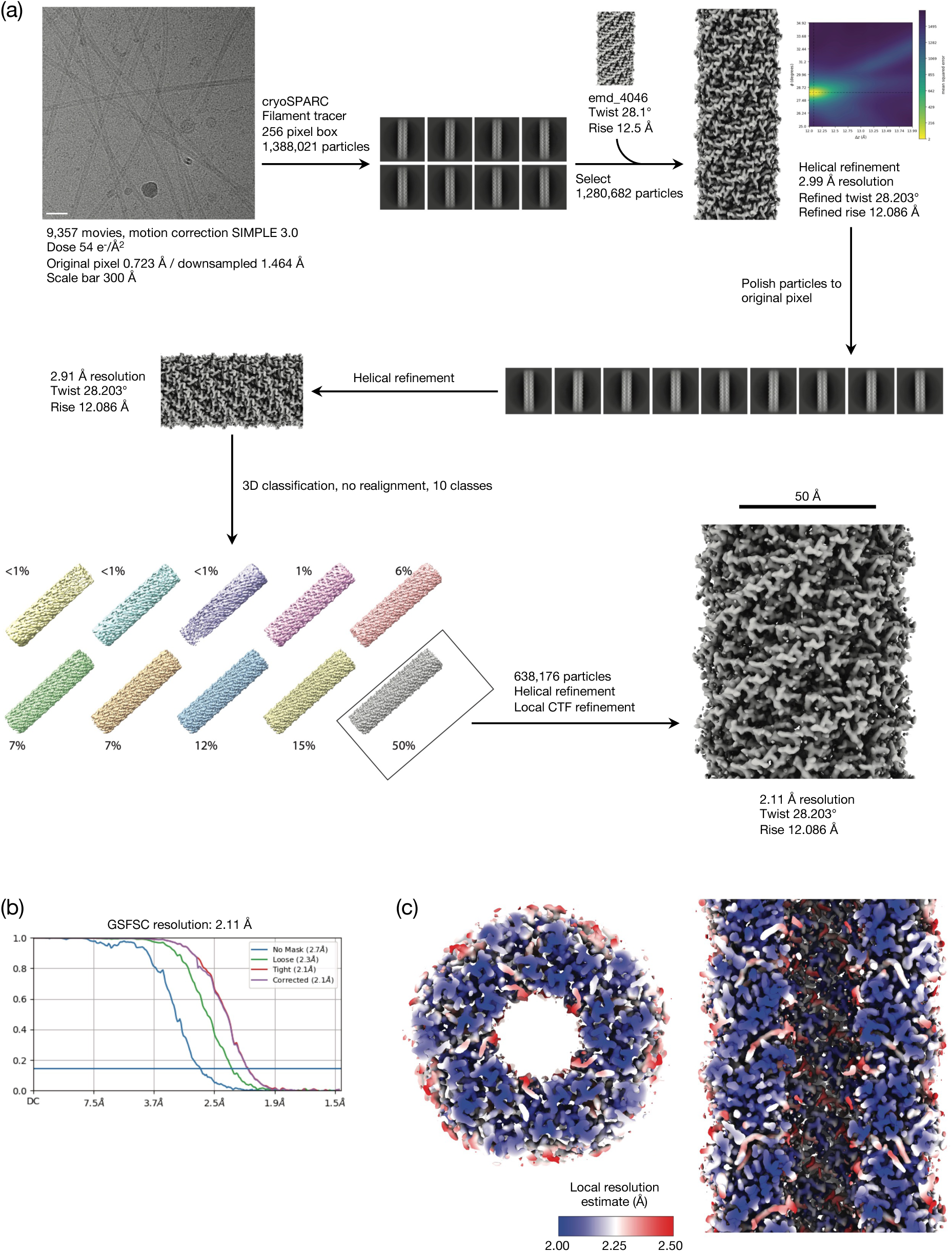
Cryo-EM workflow for the pED208 pilus. **a**, Image processing workflow, Micrograph scale bar is 300 Å. **b**, Gold-standard Fourier Shell Correlation (FSC) curves used for global resolution estimation. **c,** Local resolution estimate of the volume.

**Extended Data Figure 4.**
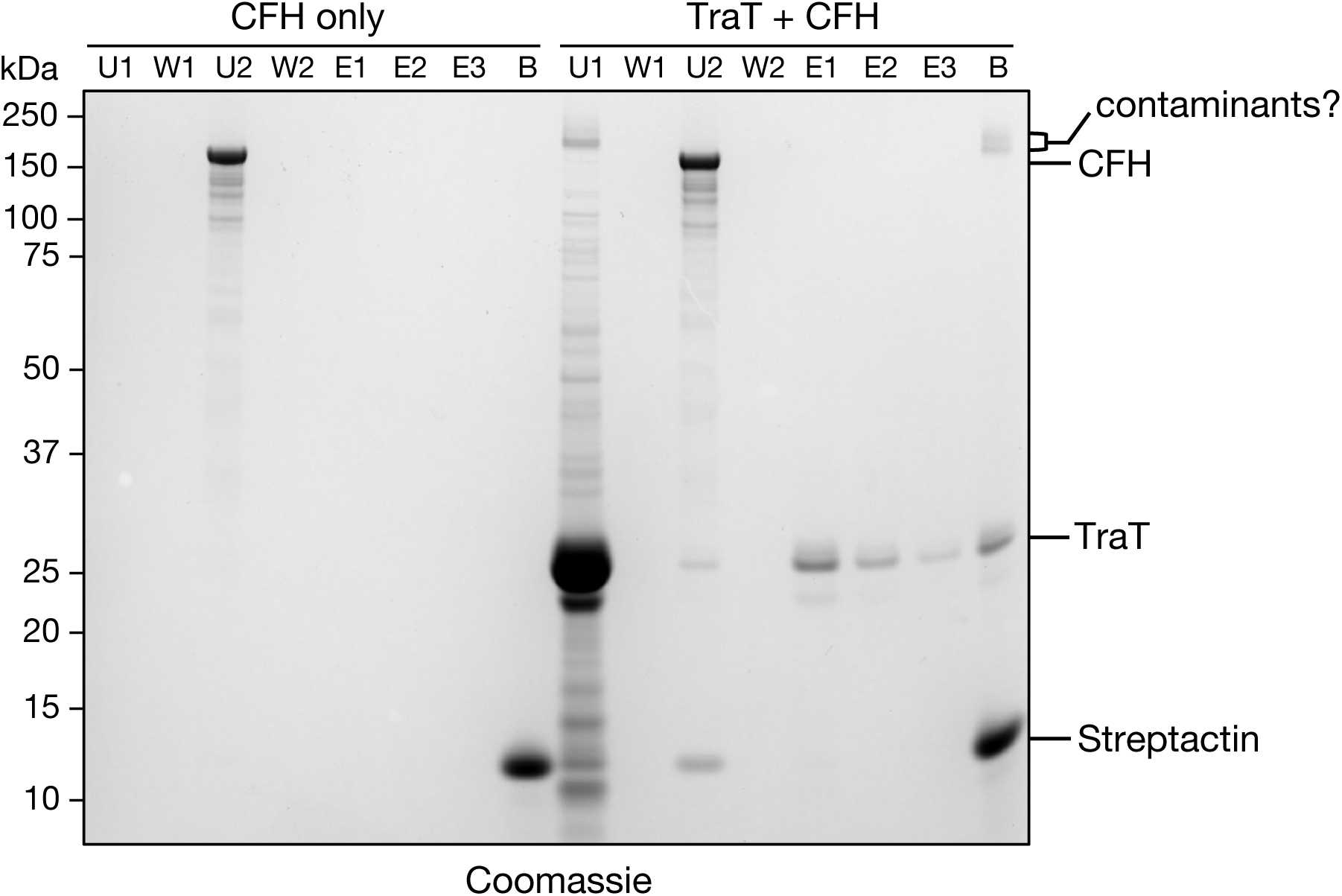
TraT fails to interact with CFH. MagStrep beads were incubated with a saturating level of purified TraT-TS (‘TraT + CFH’ lanes) or with a buffer blank (‘CFH only’ lanes). The beads were then recovered (U1 is the residual unbound fraction), washed (W1 is the final wash fraction), and incubated with mouse CFH. The beads were then recovered (U2 is the residual unbound fraction), washed (W2 is the final wash fraction), subjected to elution with desthiobiotin (elution fractions E1-E3), before being boiled at 100°C for 5 min in Laemmli buffer (B sample). Fractions were analyzed on a Coomassie Blue stained SDS-PAGE gel.

**Extended Data Figure 5.**
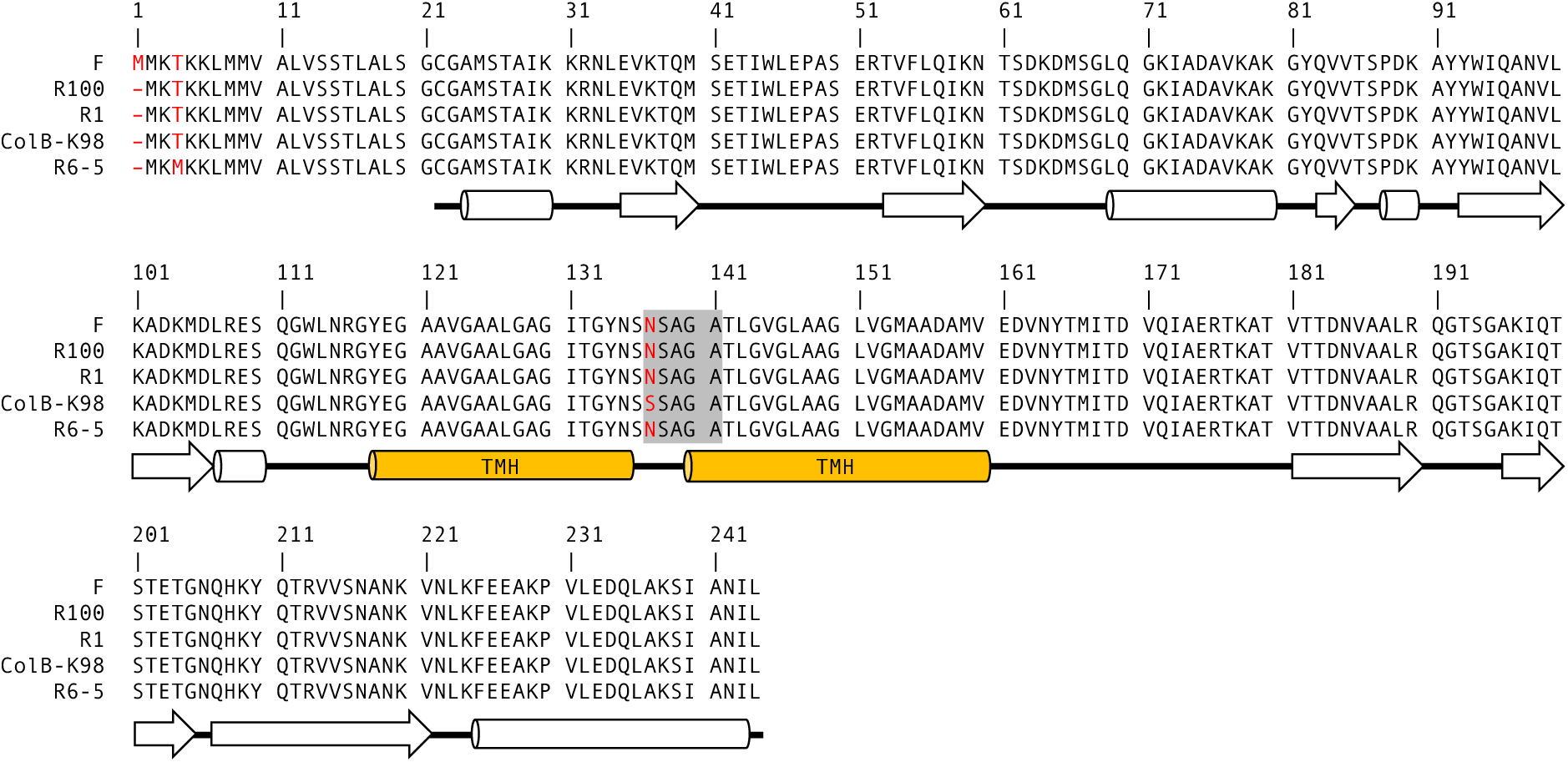
Sequence alignment of F TraT with close homologues encoded by other IncF plasmids. The alignment was performed using Multalin^67^. Sequence differences are indicated in red. The shaded region indicates the specificity region proposed in earlier work. The secondary structure diagram is plotted according to the cryo-EM structure of TraT (cylinders for α helices, arrows for β sheets). Orange cylinders represent transmembrane helices (TMH). GenBank accession numbers for the protein sequences from different plasmids are: BAA97971.1 (F), WP_000850422.1 (R100), KY749247.1 (R1), CAA32704.1 (ColB-K98), CAA36788.1 (R6-5).

## References

1 Fraikin, N., Couturier, A. & Lesterlin, C. The winding journey of conjugative plasmids toward a novel host cell. Curr Opin Microbiol 78, 102449 (2024). 10.1016/j.mib.2024.102449

2 Lederberg, J. & Tatum, E. L. Gene recombination in Escherichia coli. Nature 158, 558–558 (1946).

3 Castaneda-Barba, S., Top, E. M. & Stalder, T. Plasmids, a molecular cornerstone of antimicrobial resistance in the One Health era. Nat Rev Microbiol 22, 18–32 (2024). 10.1038/s41579-023-00926-x

4 Lederberg, J., Cavalli, L. L. & Lederberg, E. M. Sex Compatibility in Escherichia Coli. Genetics 37, 720–730 (1952). 10.1093/genetics/37.6.720

5 Virolle, C., Goldlust, K., Djermoun, S., Bigot, S. & Lesterlin, C. Plasmid transfer by conjugation in gram-negative bacteria: from the cellular to the community level. Genes 11, 1239 (2020).

6 Arutyunov, D. & Frost, L. S. F conjugation: back to the beginning. Plasmid 70, 18–32 (2013). 10.1016/j.plasmid.2013.03.010

7 Koraimann, G. Spread and Persistence of Virulence and Antibiotic Resistance Genes: A Ride on the F Plasmid Conjugation Module. EcoSal Plus 8 (2018). 10.1128/ecosalplus.ESP-0003-2018

8 Costa, T. R. D., Patkowski, J. B., Macé, K., Christie, P. J. & Waksman, G. Structural and functional diversity of type IV secretion systems. Nature Reviews Microbiology (2023). 10.1038/s41579-023-00974-3

9 Macé, K. et al. Cryo-EM structure of a type IV secretion system. Nature 607, 191–196 (2022).

10 Hu, B., Khara, P. & Christie, P. J. Structural bases for F plasmid conjugation and F pilus biogenesis in Escherichia coli. Proceedings of the National Academy of Sciences 116, 14222–14227 (2019).

11 Costa, T. R. D. et al. Structure of the Bacterial Sex F Pilus Reveals an Assembly of a Stoichiometric Protein-Phospholipid Complex. Cell 166, 1436–1444.e1410 (2016). 10.1016/j.cell.2016.08.025

12 Clarke, M., Maddera, L., Harris, R. L. & Silverman, P. M. F-pili dynamics by live-cell imaging. Proceedings of the National Academy of Sciences 105, 17978–17981 (2008). 10.1073/pnas.0806786105

13 Harrington, L. C. & Rogerson, A. C. The F pilus of Escherichia coli appears to support stable DNA transfer in the absence of wall-to-wall contact between cells. Journal of bacteriology 172, 7263–7264 (1990).

14 Goldlust, K. et al. The F pilus serves as a conduit for the DNA during conjugation between physically distant bacteria. Proceedings of the National Academy of Sciences 120, e2310842120 (2023). 10.1073/pnas.2310842120

15 Garcillán-Barcia, M. P. & de la Cruz, F. Why is entry exclusion an essential feature of conjugative plasmids? Plasmid 60, 1–18 (2008). 10.1016/j.plasmid.2008.03.002

16 Couturier, A. F., Nathan; Lesterlin, Christian. Exclusion Systems Preserve Host Cell Homeostasis and fitness, Ensuring Successful Dissemination of Conjugative Plasmids and Associated Resistance Genes. bioRxiv (2025). 10.1101/2025.04.18.649494

17 Novick, R. P. Extrachromosomal inheritance in bacteria. Bacteriological Reviews 33, 210–263 (1969).

18 Achtman, M., Kennedy, N. & Skurray, R. Cell-cell interactions in conjugating Escherichia coli: role of traT protein in surface exclusion. Proceedings of the National Academy of Sciences 74, 5104–5108 (1977).

19 Achtman, M., Manning, P. A., Kusecek, B., Schwuchow, S. & Neil, W. Genetic analysis of F sex factor cistrons needed for surface exclusion in Escherichia coli. Journal of Molecular Biology 138, 779–795 (1980). 10.1016/0022-2836(80)90065-0

20 Achtman, M., Manning, P. A., Edelbluth, C. & Herrlich, P. Export without proteolytic processing of inner and outer membrane proteins encoded by F sex factor tra cistrons in Escherichia coli minicells. Proceedings of the National Academy of Sciences 76, 4837–4841 (1979).

21 Perumal, N. B. & Minkley, E. G. The product of the F sex factor traT surface exclusion gene is a lipoprotein. Journal of Biological Chemistry 259, 5357–5360 (1984).

22 Jalajakumari, M. B. et al. Surface exclusion genes traS and traT of the F sex factor of Escherichia coli K-12: Determination of the nucleotide sequence and promoter and terminator activities. Journal of Molecular Biology 198, 1–11 (1987). 10.1016/0022-2836(87)90452-9

23 Manning, P. A., Beutin, L. & Achtman, M. Outer membrane of Escherichia coli: properties of the F sex factor traT protein which is involved in surface exclusion. Journal of Bacteriology 142, 285–294 (1980). 10.1128/jb.142.1.285-294.1980

24 Manning, P. A., Timmis, J. K., Moll, A. & Timmis, K. N. Mutants that overproduce TraTp, a plasmid-specified major outer membrane protein of Escherichia coli. Molecular and General Genetics MGG 187, 426–431 (1982). 10.1007/BF00332623

25 Li, M., Wu, M., Sun, Y. & Sun, L. Edwardsiella tarda TraT is an anti-complement factor and a cellular infection promoter. Communications Biology 5, 637 (2022). 10.1038/s42003-022-03587-3

26 Taylor, I. M., Harrison, J. L., Timmis, K. N. & O’Connor, C. D. The TraT lipoprotein as a vehicle for the transport of foreign antigenic determinants to the cell surface of Escherichia coli K12: structure–function relationships in the TraT protein. Molecular Microbiology 4, 1259–1268 (1990). 10.1111/j.1365-2958.1990.tb00705.x

27 Anthony, K. G., Sherburne, C., Sherburne, R. & Frost, L. S. The role of the pilus in recipient cell recognition during bacterial conjugation mediated by F-like plasmids. Molecular Microbiology 13, 939–953 (1994). 10.1111/j.1365-2958.1994.tb00486.x

28 Harrison, J. L., Taylor, I. M., Platt, K. & O’Connor, C. D. Surface exclusion specificity of the TraT lipoprotein is determined by single alterations in a five-amino-acid region of the protein. Molecular Microbiology 6, 2825–2832 (1992). 10.1111/j.1365-2958.1992.tb01462.x

29 Moll, A., Manning, P. A. & Timmis, K. N. Plasmid-determined resistance to serum bactericidal activity: a major outer membrane protein, the traT gene product, is responsible for plasmid-specified serum resistance in Escherichia coli. Infection and Immunity 28, 359–367 (1980). 10.1128/iai.28.2.359-367.1980

30 Binns, M. M., Mayden, J. & Levine, R. P. Further characterization of complement resistance conferred on Escherichia coli by the plasmid genes traT of R100 and iss of ColV,I-K94. *Infection and Immunity* **35**, 654-659 (1982). 10.1128/iai.35.2.654-659.1982

31 Pramoonjago, P. et al. Role of TraT protein, an anticomplementary protein produced in Escherichia coli by R100 factor, in serum resistance. The Journal of Immunology 148, 827–836 (1992). 10.4049/jimmunol.148.3.827

32 Ogata, R. T. & Levine, R. P. Characterization of complement resistance in Escherichia coli conferred by the antibiotic resistance plasmid R100. The Journal of Immunology 125, 1494 (1980).

33 Ogata, R. T., Winters, C. & Levine, R. P. Nucleotide sequence analysis of the complement resistance gene from plasmid R100. Journal of Bacteriology 151, 819–827 (1982). 10.1128/jb.151.2.819-827.1982

34 Agüero, M. E., Aron, L., DeLuca, A. G., Timmis, K. N. & Cabello, F. C. A plasmid-encoded outer membrane protein, TraT, enhances resistance of Escherichia coli to phagocytosis. Infection and Immunity 46, 740–746 (1984). 10.1128/iai.46.3.740-746.1984

35 Riede, I. & Eschbach, M.-L. Evidence that TraT interacts with OmpA of Escherichia coli. FEBS Letters 205, 241–245 (1986). 10.1016/0014-5793(86)80905-X

36 Seddon, C. et al. Cryo-EM structure and evolutionary history of the conjugation surface exclusion protein TraT. Nat Commun 16, 659 (2025). 10.1038/s41467-025-55834-w

37 Malojcic, G. et al. LptE binds to and alters the physical state of LPS to catalyze its assembly at the cell surface. Proc Natl Acad Sci U S A 111, 9467–9472 (2014). 10.1073/pnas.1402746111

38 Aly, K. A. & Baron, C. The VirB5 protein localizes to the T-pilus tips in Agrobacterium tumefaciens. Microbiology 153, 3766–3775 (2007). 10.1099/mic.0.2007/010462-0

39 Michaelis, C., Ciosk, R. & Nasmyth, K. Cohesins: Chromosomal Proteins that Prevent Premature Separation of Sister Chromatids. Cell 91, 35–45 (1997). 10.1016/S0092-8674(01)80007-6

40 Fuchs, J. r., Lorenz, A. & Loidl, J. Chromosome associations in budding yeast caused by integrated tandemly repeated transgenes. Journal of Cell Science 115, 1213–1220 (2002). 10.1242/jcs.115.6.1213

41 Dong, C. et al. Wza the translocon for E. coli capsular polysaccharides defines a new class of membrane protein. Nature 444, 226–229 (2006). 10.1038/nature05267

42 Mace, K. et al. Cryo-EM structure of a type IV secretion system. Nature 607, 191–196 (2022). 10.1038/s41586-022-04859-y

43 Garcillan-Barcia, M. P. & de la Cruz, F. Why is entry exclusion an essential feature of conjugative plasmids? Plasmid 60, 1–18 (2008). 10.1016/j.plasmid.2008.03.002

44 Willetts, N. & Maule, J. Interactions between the surface exclusion systems of some F-like plasmids. Genetical Research 24, 81–89 (1974). 10.1017/S0016672300015093

45 Low, W. W. et al. Mating pair stabilization mediates bacterial conjugation species specificity. Nature Microbiology (2022). 10.1038/s41564-022-01146-4

46 Frankel, G. et al. Plasmids pick a bacterial partner before committing to conjugation. Nucleic Acids Res 51, 8925–8933 (2023). 10.1093/nar/gkad678

47 Klimke William, A. & Frost Laura, S. Genetic Analysis of the Role of the Transfer Gene,traN, of the F and R100–1 Plasmids in Mating Pair Stabilization during Conjugation. *Journal of Bacteriology* **180**, 4036-4043 (1998). 10.1128/JB.180.16.4036-4043.1998

48 Couves, E. C. & Bubeck, D. Capturing pore-forming intermediates of MACPF and binary toxin assemblies by cryoEM. Curr Opin Struct Biol 75, 102401 (2022). 10.1016/j.sbi.2022.102401

49 Johnson, D. A. & Willetts, N. S. Construction and characterization of multicopy plasmids containing the entire F transfer region. Plasmid 4, 292–304 (1980). 10.1016/0147-619X(80)90068-2

50 Minkley, E. G. & Ippen-Ihler, K. Identification of a membrane protein associated with expression of the surface exclusion region of the F transfer operon. Journal of Bacteriology 129, 1613–1622 (1977). 10.1128/jb.129.3.1613-1622.1977

51 Datsenko, K. A. & Wanner, B. L. One-step inactivation of chromosomal genes in Escherichia coli K-12 using PCR products. Proceedings of the National Academy of Sciences 97, 6640–6645 (2000).

52 Granato, E. T., Smith, W. P. J. & Foster, K. R. Collective protection against the type VI secretion system in bacteria. The ISME Journal 17, 1052–1062 (2023). 10.1038/s41396-023-01401-4

53 Merlin, C., McAteer, S. & Masters, M. Tools for characterization of Escherichia coli genes of unknown function. J Bacteriol 184, 4573–4581 (2002). 10.1128/jb.184.16.4573-4581.2002

54 Miroux, B. & Walker, J. E. Over-production of Proteins in Escherichia coli: Mutant Hosts that Allow Synthesis of some Membrane Proteins and Globular Proteins at High Levels. Journal of Molecular Biology 260, 289–298 (1996). 10.1006/jmbi.1996.0399

55 Bertani, G. Studies on lysogenesis I: the mode of phage liberation by lysogenic Escherichia coli. Journal of bacteriology 62, 293–300 (1951).

56 Caesar, J. et al. SIMPLE 3.0. Stream single-particle cryo-EM analysis in real time. J Struct Biol X 4, 100040 (2020). 10.1016/j.yjsbx.2020.100040

57 Punjani, A., Zhang, H. & Fleet, D. J. Non-uniform refinement: adaptive regularization improves single-particle cryo-EM reconstruction. Nat Methods 17, 1214–1221 (2020). 10.1038/s41592-020-00990-8

58 Williams, C. J. et al. MolProbity: More and better reference data for improved all-atom structure validation. Protein Sci 27, 293–315 (2018). 10.1002/pro.3330

59 Jumper, J. et al. Highly accurate protein structure prediction with AlphaFold. Nature (2021). 10.1038/s41586-021-03819-2

60 Pettersen, E. F. et al. UCSF ChimeraX: Structure visualization for researchers, educators, and developers. Protein Sci 30, 70–82 (2021). 10.1002/pro.3943

61 Afonine, P. V. et al. Real-space refinement in PHENIX for cryo-EM and crystallography. Acta Crystallogr D Struct Biol 74, 531–544 (2018). 10.1107/S2059798318006551

62 Cutler, K. J. et al. Omnipose: a high-precision morphology-independent solution for bacterial cell segmentation. Nat Methods 19, 1438–1448 (2022). 10.1038/s41592-022-01639-4

63 Schnitzbauer, J., Strauss, M. T., Schlichthaerle, T., Schueder, F. & Jungmann, R. Super-resolution microscopy with DNA-PAINT. Nature Protocols 12, 1198–1228 (2017). 10.1038/nprot.2017.024

64 Glaser, F. et al. ConSurf: identification of functional regions in proteins by surface-mapping of phylogenetic information. Bioinformatics 19, 163–164 (2003). 10.1093/bioinformatics/19.1.163

65 Suzek, B. E. et al. UniRef clusters: a comprehensive and scalable alternative for improving sequence similarity searches. Bioinformatics 31, 926–932 (2015). 10.1093/bioinformatics/btu739

66 Lomize, M. A., Pogozheva, I. D., Joo, H., Mosberg, H. I. & Lomize, A. L. OPM database and PPM web server: resources for positioning of proteins in membranes. Nucleic Acids Research 40, D370–D376 (2012). 10.1093/nar/gkr703

67 Corpet, F. Multiple sequence alignment with hierarchical clustering. Nucleic Acids Research 16, 10881–10890 (1988). 10.1093/nar/16.22.10881

